# Modulation of metabolic functions through Cas13d-mediated gene knockdown in liver

**DOI:** 10.1101/2020.02.17.945014

**Authors:** Bingbing He, Wenbo Peng, Jia Huang, Hang Zhang, Yingsi Zhou, Xiali Yang, Jing Liu, Zhijie Li, Chunlong Xu, Mingxing Xue, Hui Yang, Pengyu Huang

## Abstract

RNA knockdown *in vivo* carries significant potential for disease modelings and therapies. Despite the emerging approaches of CRISPR/Cas9-mediated permanent knock out of targeted genes, strategies targeting RNA for disruption are advantageous in the treatment of acquired metabolic disorders when permanent modification of the genome DNA is not appropriate, and RNA virus infection diseases when pathogenic DNA is not available (such as SARS-Cov-2 and MERS infections). Recently, Cas13d, a family of RNA-targeting CRISPR effectors, has been shown to accomplish robust down-regulation of cellular RNAs in mammalian cells *in vitro*. Among the various Cas13d subtypes, CasRx (RfxCas13d) showed the most potent RNA knockdown efficiency in HEK293T cells. However, the RNA-targeting activity of Cas13d still needs to be verified *in vivo*. In this study, the CasRx system was demonstrated to efficiently and functionally knock down genes related to metabolism functions, including *Pten, Pcsk9* and *lncLstr*, in mouse hepatocytes. CasRx-mediated simultaneous knockdown of multiple genes was also achieved by sgRNA arrays, providing a useful strategy to modulate complex metabolism networks. Moreover, the AAV (adeno-associated virus)-mediated delivery of *CasRx* and *Pcsk9* sgRNAs into mouse liver successfully decreased serum PCSK9, resulting in significant reduction of serum cholesterol levels. Importantly, CasRx-mediated knockdown of *Pcsk9* is reversible and *Pcsk9* could be repeatedly down-regulated, providing an effective strategy to reversibly modulate metabolic genes. The present work supplies a successful proof-of-concept trial that suggests efficient and regulatory knockdown of target metabolic genes for a designed metabolism modulation in the liver.

## Introduction

Targeted inhibition of a metabolism regulatory gene is often used for modeling and developing therapies of metabolic diseases (Moller, 2012; Saviano and Baumert, 2019). In the past decades, many strategies of metabolic regulations were achieved through using various modulators, including numerous small molecular compounds, nucleic acids, and therapeutic polypeptides/proteins targeting individual or multiple defined molecular products, such as enzymes, circulating proteins, cell-surface receptors and cellular RNAs (Moller, 2012). Applications of metabolic modulation have great potential for disease modeling and therapies. However, developing novel approaches with more specific and more flexible modulation of metabolic genes *in vivo* is still challenging, because natural and synthesized modulators with high targeting specificity are theoretically limited.

In recent years, an increasing number of genetic modification tools has emerged. Representing one of the greatest breakthroughs, the CRISPR/Cas system offers sequence-specific DNA editing methods to correct mutant genes in inherited metabolic diseases, and shows remarkable benefits to the establishments of metabolic disease animal models, such as inherited tyrosinemia (Barrangou and Doudna, 2016; Rossidis et al., 2018; Xue et al., 2014; Yao et al., 2017). However, permanent modification of DNA is usually not an optimal strategy for the therapies of acquired metabolic disorders.

Recent breakthroughs with RNA-guided RNA-targeting systems, such as class 2 type VI CRISPR-Cas effectors, were reported for the remarkable capability of RNA editing without permanent modification of DNA. Theoretically, RNA-targeting systems could provide a much safer approach for gene silencing (Abudayyeh et al., 2017; Cox et al., 2017; Konermann et al., 2018; Pineda et al., 2019). Among such systems, CasRx was recently found to be advantageous, mostly because it is more efficient and has more robust activation during RNA-guided RNA cleavage in mammalian cells *in vitro* (Konermann et al., 2018; Yan et al., 2018). CasRx is a type VI-D effector (Cas13d) with a small size that can be packaged into AAVs, and thus has great potential for translational medicine (Konermann et al., 2018; Yan et al., 2018; Zhang et al., 2018). In this study, we set out to explore the feasibility of using the CasRx system for the targeted silencing of metabolic genes.

## Results

As one of the metabolism regulatory genes, *Pten* was chosen first in our study to investigate whether CasRx could generally target metabolic genes for the efficient knockdown. 10 sgRNAs that were designed to target to the coding sequence of *Pten* mRNAs were prepared (Fig. S1A). *CasRx* and each *Pten* sgRNA were introduced through plasmids transfection into mouse neuroblastoma (N2a) cells (Fig. S2A). After 48 hours, the N2a cells with the transfected CasRx and each *Pten* sgRNA were analyzed to evaluate their levels of *Pten* mRNA. As expected, all of our designed *Pten* sgRNAs revealed successful down-regulation of the level of *Pten* mRNA (Fig. S2A).

Next, improvement of knockdown efficiency was tested on the strategy to combine the different *Pten* sgRNA individuals (Fig. S2B, C, D). After sgPten-5 and sgPten-6 were combined in treatment, the reduction of *Pten* mRNA levels successfully reached 16±3%, the most efficient of all combinations (Fig. 1A, B and Fig. S2D). Previously, *Pten* was known as a metabolic regulator that suppressed the insulin signaling by inhibiting the PI3K/AKT pathway (Cho et al., 2001; Stiles et al., 2004). Deletion of *Pten* promoted AKT phosphorylation (Stiles et al., 2004). Accordingly, such effects were also shown in our results after the successful knockdown of *Pten* through the combination of sgPten-5 and sgPten-6. In the N2a cells that were transfected with CasRx, sgPten-5 and sgPten-6, the PTEN protein levels were significanty decreased while the AKT phosphorylation levels were significantly elevated (Fig. 1C).

**Figure 1:**
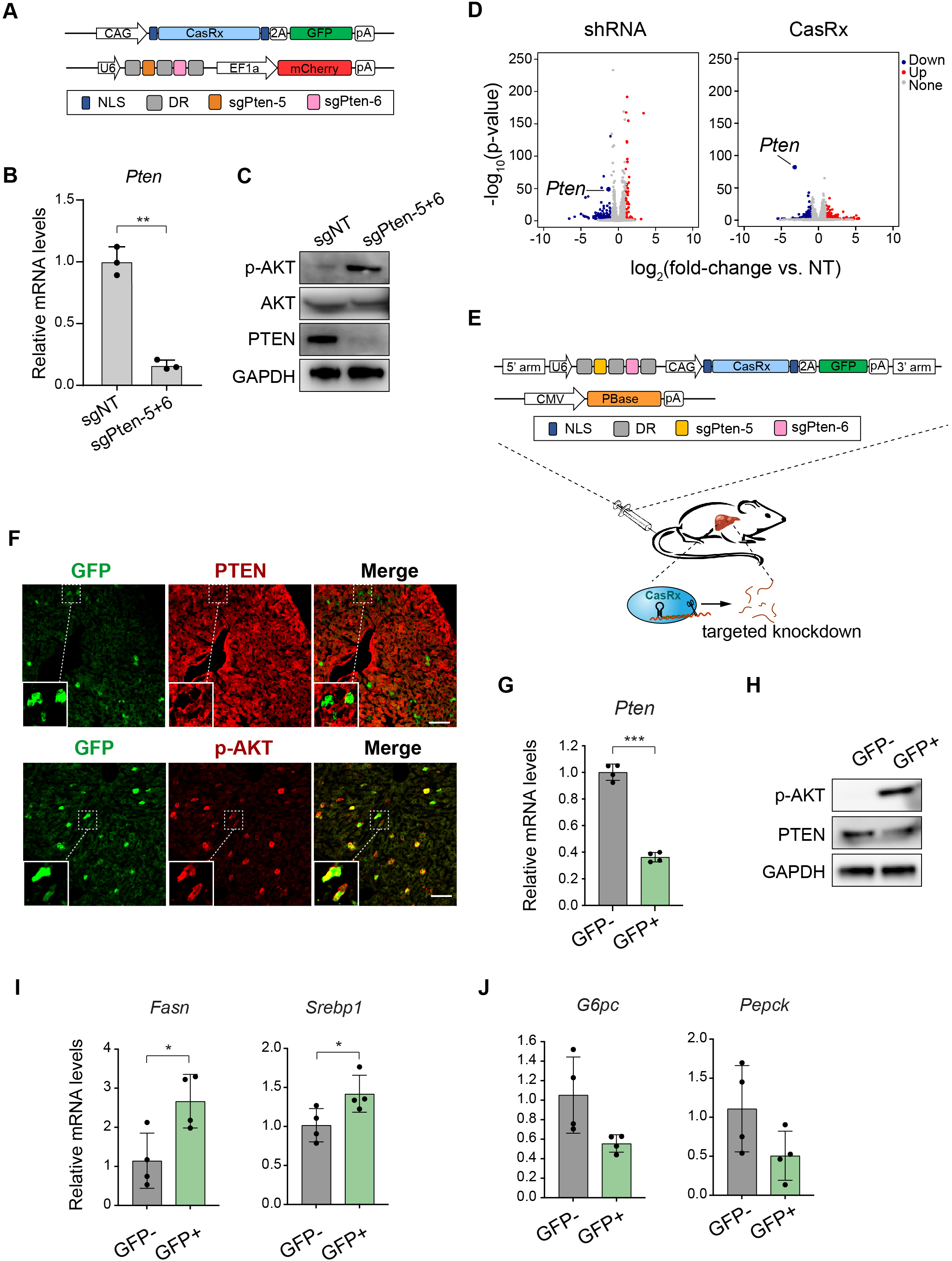
CasRx-mediated knockdown of *Pten in vitro* and *in vivo*. **A.** Schematic of plasmids used for CasRx-mediated *Pten* knockdown in N2a cells with the combination of sgPten-5 and sgPten-6. **B.** Knockdown of *Pten* by a combination of sgPten-5 and sgPten-6 in N2a cells (n=3). **C.** PTEN and its downstream signaling protein AKT were analyzed by western blot in N2a cells transfected with plasmids encoding CasRx and two sgRNAs. **D.** Left, Volcano plot of differential transcript levels between Pten-targeting (sgPten-5+6) and non-targeting (NT) CasRx guide RNAs (n=3). Right, Volcano plot of differential transcript levels between Pten-targeting (shPten-5+6, n=2) and non-targeting (NT) CasRx guide RNAs (n=3). **E.** Design of PBase and CasRx plasmids used for knockdown of *Pten* in hepatocytes by hydrodynamic tail-vein injection. **F.** Representative PTEN and phospho-Akt (Ser473) immunofluorescence staining on liver sections from sgPten treated mice (n=3). The lower-left insets are high-magnification views, indicating hepatocytes with reduced PTEN staining and corresponding improved phospho-Akt staining. Scale bar: 100 μm. **G, H.** GFP+ hepatocytes were sorted for the quantification of *Pten* mRNA levels (n=4) (**G**) and protein levels (**H**). Elevated phosphorylation of AKT was observed in GFP+ hepatocytes (**H**). **I.** The expressions of fatty acid synthesis-associated genes, *Fasn* and *Srebp1*, quantified by qPCR (n=4). **J.** The expressions of glucose metabolism genes, *G6pc* and *Pepck* quantified by qPCR (n=4). Data are represented as mean with SD. *p<0.05, ***p<0.001.

Unintended wide-spread off-target silencing is one of the major hurdles for the applications of RNA interference (RNAi) (Jackson et al., 2003; Pecot et al., 2011). The CasRx system has, however, been reported to be with significantly reduced off-target events compared to other RNAi approaches (Konermann et al., 2018). Here, we set out to compare the CasRx system with spacer sequence-match shRNAs in knockdown of *Pten* via transient transfection in N2a cells. Consistent with previous studies, *Pten*-targeting shRNAs rendered wide-spread off-target transcript changes (Fig. 1D). In contrast, transfection of the spacer-matched CasRx system induced significantly reduced transcript changes (Fig. 1D). Importantly, *Pten* was the most significantly changed gene (Fig. 1D). These results further validated the high specificity of CasRx-mediated transcript interference.

Next, CasRx-mediated knockdown of a targeted gene was broadly studied among metabolism regulatory genes. Proprotein convertase subtilisin/kexin type 9 (PCSK9) is a crucial protein of serum LDL cholesterol regulation (Lambert et al., 2012). Here, a total of 11 candidate sgRNAs were designed and tested, with the goal of targeting the coding region of *Pcsk9* (Fig. S1B). As expected, all *Pcsk9* sgRNA candidates showed the effects of robust knockdown of *Pcsk9* mRNAs in the 293T cells transfected with each sgRNA for *Pcsk9* (Fig. S2E). Moreover, CasRx-mediated targeted knockdown could also be applied for *lncLstr*, a liver-enriched lncRNA (Fig. S1C and S2F). Together, the CasRx system was able to efficiently and functionally knock down the metabolic genes, including the lncRNA gene that could be related to metabolic function, in the *in vitro* cultured mammalian cells.

After the successful establishment of a method for CasRx-mediated gene knockdown in the cultured mammalian cells *in vitro*, the feasibility of *in vivo* metabolic gene targeting was studied. The three metabolic genes *Pten, Pcsk9* and *lncLstr*, were further studied to determine whether gene knockdown was effective in hepatocytes. At first, the effect of CasRx-mediated *Pten* gene knockdown in the liver was studied through the evaluation of *Pten* mRNA levels. CasRx, sgPten-5, and sgPten-6 were delivered to the mouse liver by hydrodynamic tail-vein injection of indicated plasmids (Fig. S3A). Hydrodynamic tail-vein injection, a simple yet effective method, can be used to deliver naked DNA into hepatocytes of mouse liver *in vivo* (Kim and Ahituv, 2013). After 96 hours of plasmid injection, individual hepatocytes were dissociated through liver perfusion. Hepatocytes expressing both GFP and mCherry implied successful vector delivery and were purified through cell sorting. Though not mathematically significant, results suggested that the *Pten* mRNA level in GFP+/mCherry+ hepatocytes had decreased in comparison with that in GFP-/mCherry-hepatocytes (Fig. S3B).

Second, the *in vivo* CasRx-mediated *Pcsk9* gene knockdown in mouse liver was studied. After hydrodynamic injection of the CasRx system with sgPcsk9-5 and sgPcsk9-6, *Pcsk9* mRNA levels in GFP+/mCherry+ hepatocytes were significantly decreased to 28.8±10.5% of that in GFP-/mCherry-hepatocytes (Fig. S3C), which also significantly reduced PCSK9 protein level in GFP+/mCherry+ hepatocytes (Fig. S3D).

Next, the feasibility of the CasRx system in knockdown of long non-coding RNAs (lncRNAs) *in vivo* was investigated. *LncLstr* is a liver-enriched lncRNA, which regulates lipid metabolism by inhibition of *Cyp8b1* (Li et al., 2015). Similar with the above methods for *in vivo* vector deliveries, hydrodynamic injection of plasmids encoding CasRx and those encoding both sglncLstr-5 and sglncLstr-6 were performed (Fig. S3A). Hepatocytes that received these two plasmids showed significant reduction of *lncLstr* (Fig. S3E). CasRx-mediated *lncLstr* knockdown was sufficient to result in reduced expression of *Cyp8b1* (Fig. S3F). Thus, efficient knockdown of lncRNA was successfully obtained in hepatocytes using the CasRx system.

In an effort to explore the feasibility of using the CasRx system in simultaneous knockdown of multiple genes, we cloned a vector encoding CasRx and GFP together with 6 sgRNAs targeting *Pten, Pcsk9* and *lncLstr* (Fig. S3G). Hydrodynamic tail-vein injection of this plasmid into the liver simultaneously decreased all three targeted genes, providing a powerful platform for simultaneous knockdown of multiple metabolic genes (Fig. S3G).

To evaluate the functional modulation of metabolic genes by CasRx, we first employed the piggyBac transposon system to transfer CasRx and *Pten*-targeting sgRNAs to the hepatocytes by hydrodynamic injection (Fig. 1E). Four days after injection of GFP-expressing CasRx plasmids, we observed scattered hepatocytes with reduced PTEN staining (Fig. 1F). Importantly, GFP+ hepatocytes were also negative or low for PTEN staining (Fig. 1F). We also observed increased AKT phosphorylation in hepatocytes receiving the CasRx plasmids, suggesting a functional knockdown of *Pten* in these cells (Fig. 1F). To further characterize the hepatocytes receiving CasRx plasmids, we isolated GFP+ hepatocytes by FACS sorting. The GFP+ hepatocytes showed significantly reduced expression of *Pten* and increased AKT phosphorylation (Fig. 1G, H).

To explore the functional activation of AKT signaling cascades, we investigated the response of lipid and glucose metabolism-related genes downstream of AKT. The results showed that *Pten* knockdown significantly increased the expressions of *Fasn* (fatty acid synthase) and *Srebp1* (sterol regulatory element-binding protein-1) in the GFP+ hepatocytes (Fig. 1I), which was consistent with previous findings (Stiles et al., 2004). Pten-inhibited PI3K/AKT signaling was found to be critical for lipogenesis regulation in the liver, while the activated AKT repressed SREBP1, a key transcription factor required for the expression of *Fasn* (Haeusler et al., 2014; He et al., 2010). Therefore, decrease of PTEN promoted the expression of *Fasn* and *Srebp1* (Fig. 1I). *Pten* is also involved in regulation of glucose metabolism in the liver (Gross et al., 2009). AKT positively regulates the transcriptions of glucose-6-phosphatase (*G6pc*) and phosphoenolpyruvate carboxykinase (*Pepck*) in hepatocytes (Gross et al., 2009). Here, the CasRx-mediated knockdown of *Pten*, a negative regulator of AKT activity, led to the decreased expression of *G6pc* and *Pepck* in GFP+ hepatocytes (Fig. 1J). Together, the above results indicated that the CasRx system could be used to functionally knock down *Pten in vivo*.

Next, AAV8, an efficient liver-targeted gene delivery system, was applied to further increase the effect of *in vivo Pcsk9* knockdown (Fig. 2A) (Shen et al., 2007). PCSK9 is secreted by hepatocytes, and has great promise as a candidate of drug targets among all regulators of serum cholesterol (Steinberg and Witztum, 2009). Accumulating evidence shows that inhibition of PCSK9 lowered serum cholesterol levels (Chan et al., 2009; Rossidis et al., 2018). Increased plasma LDL cholesterol level is one of the major causes for coronary heart disease (CHD) and cardiovascular disease (CVD), as well as many other diseases (LaRosa et al., 1990; Law et al., 1994). Thus, we chose the CasRx system applied to *Pcsk9* gene knockdown in hepatocytes for the purpose of reducing serum cholesterol levels.

**Figure 2:**
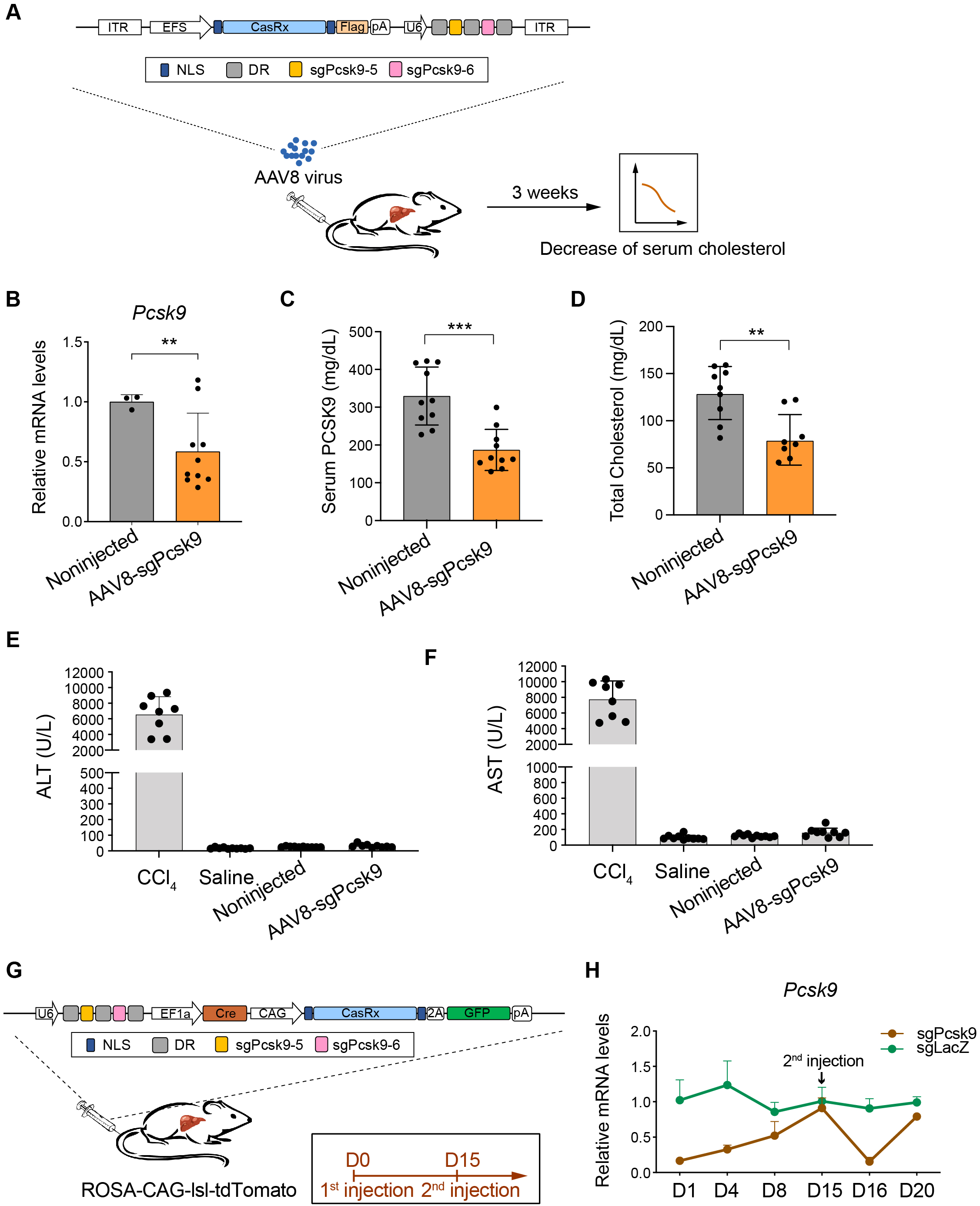
Reduction of serum cholesterol and reversible modulation of *Pcsk9* by CasRx-mediated knockdown of *Pcsk9* in the liver. **A.** Experimental scheme. **B.** Quantification of *Pcsk9* mRNA levels of livers from AAV8-injected (n=10) and noninjected mice (n=3). **C.** Serum PCSK9 protein levels were quantified at 3 weeks (n=10). **D.** Quantification of serum total cholesterol levels at 3 weeks (sgNT, n=9; sgPcsk9, n=8). **E, F.** Serum ALT and AST were quantified in CCl_4_-injected (n=8), saline-injected (n=10), Noninjected (n=10) and AAV8-sgPcsk9-injected (n=9) mice. Data are represented as mean ± SD. **p<0.01, ***p<0.001. **G.** Plasmids expressing CasRx, sgRNAs, Cre and GFP were delivered to ROSA-CAG-lsl-tdTomato mouse livers by hydrodynamic tail-vein injection. The first and second injections were given at day 0 and day 15, respectively. **H.** Quantification of *Pcsk9* mRNA levels of hepatocytes receiving plasmids. For D1, D4, D8, D16 and D20, FACS-sorted hepatocytes expressing GFP and tdTomato were used for quantification (sgPcsk9: D1, n=4; D4, D8, n=3, sgLacZ: D1, D4, D8, D16, n=4; D20, n=3). For D15, FACS-sorted GFP-/tdTomato+ hepatocytes were used for quantification (sgPcsk9: n=3, sgLacZ: n=4). Pcsk9 mRNA levels in GFP-/tdTomato-hepatocytes were used as references. Data are represented as mean ± SD.

We delivered CasRx, sgPcsk9-5 and sgPcsk9-6 to the liver by AAV8, which significantly reduced *Pcsk9* mRNA level (Fig. 2A, B). The serum PCSK9 levels in *Pcsk9* knockdown mice was reduced to 56.8±15.6% of those in non-injected wild-type mice 3 weeks after AAV infection (Fig. 2C). Serum total cholesterol levels were reduced to 61.6±19.4% of normal levels (p=0.002) (Fig. 2D). Moreover, the liver function of AAV-injected mice, saline-injected and non-injected mice were similar, and we did not observe obvious liver injuries in these mice (Fig. 2E, F). Thus, the CasRx system provides an efficient tool targeting *Pcsk9* to reduce serum cholesterol *in vivo*.

To investigate whether CasRx-mediated gene knockdown is reversible after removal of the CasRx system, we delivered CasRx plasmids expressing GFP, Cre and Pcsk9-targeting sgRNAs to the livers of ROSA-CAG-lsl-tdTomato mice by hydrodynamic injection (Fig. 2G). CasRx plasmids expressing LacZ-targeting sgRNA were used as controls. Hepatocytes receiving the plasmids could be traced by the expression of red fluorescent protein tdTomato. One day after injection of plasmids, *Pcsk9* mRNA levels in hepatocytes receiving CasRx plasmids (tdTomato+) decreased to 16.5±4.9% of those in hepatocytes without plasmids (GFP-/tdTomato-) (Fig. 2H). Along with the gradual loss of CasRx plasmids in hepatocytes, the *Pcsk9* mRNA levels gradually recovered to normal levels 15 days after injection (Fig. 2H and Fig. S4A). Importantly, a second-round injection of the plasmids achieved a similar efficiency of *Pcsk9* knockdown compared to that in the first-round of plasmids injection (Fig. 2H). These results suggested a remarkable superiority of the CasRx system for its reversibility of gene knockdown.

## Discussion

Through these early studies we have successfully obtained firsthand information and experiences for realizing the potential and feasibility of the CasRx system in RNA knockdown *in vivo*. These results support our long-term goal to model and develop therapies of metabolic diseases. Over the past few years, advancements in CRISPR systems have provided plentiful tools for genome editing, epigenetic modification and transcript activation/inhibition both *in vitro* and *in vivo*. Thus, the development of a CasRx-mediated *in vivo* gene knockdown system is becoming more relied upon for disease modeling, genetic screening, mechanism studies, and therapeutic purposes. Remarkably, different from CRISPR/Cas-mediated genome editing with permanent disruption of DNA *in vivo*, CasRx-mediated modification is used to target RNAs, instead of genomic DNA (Zhang et al., 2018). This impermanent modification makes the CasRx system more valuable for modeling metabolism disorders considering the required reversible downregulation of metabolism genes. Moreover, CasRx-mediated modification can ideally also be used for therapeutic applications in clinical metabolism disorders because of its advantages in terms of reversible manipulations of gene expressions.

In this study, hepatocytes were treated as the targeting locations because they are one of the most important cell types to maintain metabolism homeostasis in the human body. Indeed, many drug candidates were first designed to target proteins that are produced by the hepatocytes in order to correct metabolic disorders. Because hepatocytes are composed of a large fraction of polyploid cells (Duncan et al., 2010), the multiple copies of an individual gene in one hepatocyte make CRISPR/Cas-mediated gene knockout less efficient. Thus, posttranscriptional silencing approaches attracted much attention for disrupting gene expression in the liver, which can rely on the CasRx system for modulation of mRNAs at the post-transcriptional level. The successful CasRx-mediated gene knockdown in hepatocytes is expected to be beneficial to similar approaches in other cells. Compared to other approaches of gene knockdown with RNA interference, the CasRx system-derived approach was more specific and efficient (Konermann et al., 2018; Yan et al., 2018). In this study, we demonstrated that the CasRx system could efficiently target RNA for knockdown in hepatocytes, providing a robust method to inactivate genes in polyploid cells *in vivo*.

Another advantage of CasRx is its small size, when considering therapeutic potential (Barrangou and Doudna, 2016; Konermann et al., 2018). Currently, CasRx is the smallest class 2 CRISPR effector available in mammalian cells (Konermann et al., 2018). The small size of 966 aa for one CasRx protein molecule makes it possible to package CasRx into AAV together with multiple guide RNAs (Konermann et al., 2018; Zhang et al., 2018) (Fig. 2A), highlighting its potential in therapeutic uses.

As an RNA-targeting CRISPR effector, CasRx is especially appealing for its potential in the treatment of RNA virus infections. A Cas9-based DNA-targeting strategy has been reported to function in disruption of genome DNA or DNA intermediates of viruses (Liu et al., 2015; Ophinni et al., 2018; Price et al., 2015; Yin et al., 2017). However, RNA viruses with neither genome DNA nor DNA intermediates, such as deadly SARS-Cov-2 (2019-nCov), SARS-Cov, MERS-Cov and influenza A virus, could not be targeted by DNA-targeting CRISPR effectors. Recently, RNA-targeting Cas13 was experimentally validated for its antiviral activity in cell lines infected with single-stranded RNA viruses (Freije et al., 2019). In this study, we validated that CasRx, a Cas13 family protein, could also functionally target RNA *in vivo*, further supporting the potential use of CasRx in RNA-targeting therapies.

## FOOTNOTES

We thank all staff of the molecular and cell biology core facility and the core imaging facility of the School of Life Science and Technology at ShanghaiTech University for their technical support. We also thank FACS facility SQ, HW and LQ in ION. P.H. is funded by the Ministry of Science and Technology of China (MOST; 2019YFA0801501, 2016YFA0100500), NSFC grants (31970687, 31571509, 31522038). H.Y. is funded by R&D Program of China (2018YFC2000100 and 2017YFC1001302), CAS Strategic Priority Research Program (XDB32060000), National Natural Science Foundation of China (31871502 and 31522037), Shanghai Municipal Science and Technology Major Project (2018SHZDZX05), and Shanghai City Committee of Science and Technology project (18411953700 and 18JC1410100).

The authors declare no competing interests.

The use and care of animals complied with the guideline of the Biomedical Research Ethics Committee of Shanghai Institutes for Biological Science, Chinese Academy of Sciences.

## Author information

Bingbing He, Wenbo Peng, Jia Huang contributed equally to this study.

### Contributions

P.H. and H.Y. designed and supervised the project. B.H. constructed the AAV plasmids, prepared virus, and performed *in vitro* cell line experiments and analysis. W.P. designed and conducted the animal experiments, tissue staining, and imaging. J.H. designed sgRNAs. J.H., C.X., X.Y., J.L. and M.X. constructed and prepared plasmids. H.Z. performed western blot experiments. B.H. and C.X. performed qPCR analysis. Z.L. performed the CCl_4_-induced liver injury experiments. Y.Z. performed the RNA-seq analysis. P.H. prepared the figures and wrote the manuscript. All authors revised and approved the manuscript.

### Corresponding authors

Correspondence to Pengyu Huang or Hui Yang.

## Electronic supplementary material

Supplementary materials

