## Supplementary Material for "Modulation of metabolic functions through Cas13d-mediated gene knockdown in liver"

**Materials and methods**

**Mice**

8 weeks old male C57BL/6 (SLAC laboratory) mice and male heterozygous Ai9 (Rosa-CAG-LSL-tdTomato-WPRE, Jackson Laboratory) mice were used for Hydrodynamic tail vein injection. The use and care of animals complied with the guideline of the Biomedical Research Ethics Committee of Shanghai Institutes for Biological Science, Chinese Academy of Sciences.

**Generation of plasmids**

The CasRx and sgRNA backbone sequences were synthesized by the HuaGene Company. Then we generated vectors: *CAG-CasRx-p2A-GFP*, *U6-BbsI-DRs-EF1α-mCherry* and *U6-BbsI-CMV-GFP.* The sgRNA sequences were inserted by BbsI restriction enzyme*.* To insert two sgRNA sequences, overlapped PCR were performed using primers in the table S2. Vectors *U6-BbsI-DRs-EF1α-mCherry-2A-Pcsk9* and *U6-BbsI-DRs-EF1α-mCherry-2A-lncLstr* were constructed for screen of sgPcsk9 and sglncLstr. Vector *PBL-U6-DR-sgPten*s*-DR-CAG-CasRx-p2A-GFP-PBR and CMV-PBase* were constructed for piggyBac-mediated transfer of CasRx and sgPtens to hepatocytes. Vector *ITR-EFS-CasRx-Flag-polyA-U6-DR-sgRNA*s*-DR-ITR* was constructed to generate AAV8. All of the primers were listed in the table S2.

**Cell culture and transfection**

N2a (Neuro-2a) and 293T cell lines were purchased from Stem Cell Bank, Chinese Academy of Sciences. N2a and 293T cell lines were cultured with DMEM (Gibco) supplemented with 10% fetal bovine serum (Gibco), 1% penicillin/streptomycin (Thermo Fisher Scientific) and 0.1 mM non-essential amino acids (Gibco) in an incubator at 37 °C with 5% CO_2_. When cells reached 90% confluence, 293T and N2a cells were passaged at a ratio of 1:6 to 24-well plates. After 12 hours, 1 μg/well plasmids were introduced into cells with Lipofectamine 3000 (Thermo Fisher Scientific) using the standard protocol. 48 hours after transfection, GFP and mCherry positive cells were sorted by BD FACS Aria II for RNA extraction. For transcriptome sequencing, 30ug plasmids expressing shRNA or CasRx with sgRNAs were transfected into 10cm dishes. Then 500k positive cells were sorted out to make a pool for sequencing.

**RNA extraction and qPCR**

The total RNA was extracted by adding 500 μL Trizol (Invitrogen), 200 μL chloroform to the cells. After centrifuge at 12000 rpm for 15 min at 4 ℃，the supernatant was transferred to a 1.5 mL RNase-free tube. Then isopropanol and 75% alcohol were used to precipitate and purify the RNA. The cDNA was prepared using HiScript Q RT SuperMix for qPCR (Vazyme, Biotech) according to manufacturer’s instructions. qPCR reactions were performed with AceQ qPCR SYBR Green Master Mix (Vazyme, Biotech). All of the reagents were precooled in advance. All of the primers used for qPCR reactions were listed in the table S3.

**Preparation of AAV8**

AAV plasmids along with adenoviral helper and AAV rep gene, AAV8 cap gene were introduced to HEK293FT cells by Polythylenimine Max (PEI, Polysciences) in DMEM+10% FBS media. Viral supernatant was harvested 7 days later, purified by iodixanol density gradient purification.

**Hydrodynamic tail vein injection and hepatocyte isolation**

Male C57BL/6 (SLAC laboratory) mice or heterozygous Ai9 (Rosa-CAG-LSL-tdTomato-WPRE, Jackson Laboratory) mice at the age of 8 weeks were used for hydrodynamic tail vein injection. Mice were infected with 1×10^11^ transducing units (TU) of AAV in 100 μL PBS by intravenous injection. Vectors for hydrodynamic tail-vein injection were prepared using the Plasmid DNA purification Midi Kit (MACHEREY-NAGEL). For hydrodynamic liver injection, plasmid DNA suspended in 2 ml saline was injected via the tail vein in 5–7 s into 8-week-old male mice. Mice were sacrificed for analysis at 1, 4, 8, 15 or 16 days post-injection of plasmids or 3 weeks post-injection of AAV. Mouse primary hepatocytes were isolated by standard two-step collagenase perfusion and purified by 40% Percoll (Sigma) at low-speed centrifugation (1000 rpm, 10 min). Hepatocytes were resuspended in DMEM plus 10% fetal bovine serum (FBS) for FACS, and specific cell populations were used for RNA extraction or western blot analysis.

**Immunofluorescence staining and western blot**

Mice were sacrificed at 4 days post-injection of PBase plasmids and CasRx/sgPten plasmids. Separated liver lobes were harvested and fixed in 4% of paraformaldehyde overnight, and stored in 30% sucrose for 12h. After embedding in OCT compound (Sakura Finetek), 10μm liver sections were used for immunofluorescence staining. The following antibodies were used: anti-Pten (Cell Signaling, 9559, 1:100), anti-pAkt S473 (Cell Signaling, 4060, 1:100). For immunoblotting, proteins were separated by SDS-PAGE, then transferred onto a PVDF membrane and identified by immunoblot analysis with appropriate primary antibodies, including GAPDH antibody (Proteintech, 10494-1-AP, 1:5000), PTEN antibody (1:2000), p-AKT S473 antibody (1:2000), AKT antibody (1:2000), PCSK9 (10007185, 1:2000). HRP-labeled Goat Anti-Rabbit IgG (H+L) (1:1000) were from Beyotime Institute of Biotechnology. The protein bands were visualized with a BeyoECL Star Kit according to the manufacturer’s instructions (Beyotime Institute of Biotechnology, P0018A).

**Serum cholesterol and PCSK9 protein mensuration, serum biochemistry**

Before whole blood collection, mice were fasted for 4 hours. Then whole blood was collected at the time of euthanasia. Whole blood were stood at room temperature for 1 hour and centrifuged at 2000 g for 20 min. Then transfer the serum into new tubes for further analysis. Serum PCSK9 protein and cholesterol were measured with Mouse Proprotein Convertase9/PCSK9 Quantikine ELISA Kit (R&D Systems) and the Infinity Cholesterol Reagent (Thermo Fisher Scientific), respectively, according to the manufacturer’s instructions. In the CCl_4_ group, WT mice were intraperitoneally injected with CCl_4_ (1 mL/kg body weight; Shanghai Hushi, Shanghai, China) and sacrificed for analysis at 24 hours after injection. Mice intraperitoneally injected with the same volume of saline were sacrificed for analysis at 24 hours after saline injection. Serum parameters of liver functions, including alanine aminotransferase (ALT), aspartate transaminase (AST), were measured by automatic biochemical analyser at the Adicon Clinical Laboratories.inc (Shanghai, China).

**RNA-seq analysis**

The transcriptome libraries were sequenced using 150 bp paired-end Illumina Xten platform. After filtering the low-quality reads with SolexaQA (V3.1.7.1) ^24^, RNA-seq reads were aligned the reads to mm10 reference genome with Hisat2 (V2.0.4) ^25^. All uniquely mapped reads were used to calculate the read counts with htseq-count (v0.11.2) ^26^. DEseq2 (1.24.0) were used to calculate differentially expressed genes ^27^. Genes with fold-change>2 and FDR<0.05 were treated as differentially expressed genes.

**Statistical analysis**

All values are reported as mean ± SD. A two-tailed unpaired Student’s t-test was used and statistical significance was determined as *p < 0.05, **p < 0.01, ***p < 0.001. Graphpad Prism 8 was used for all statistical analysis.

**Data availability**

RNA-seq data are available with the SRA accession number SRP218720.

**Supplementary figure legends**

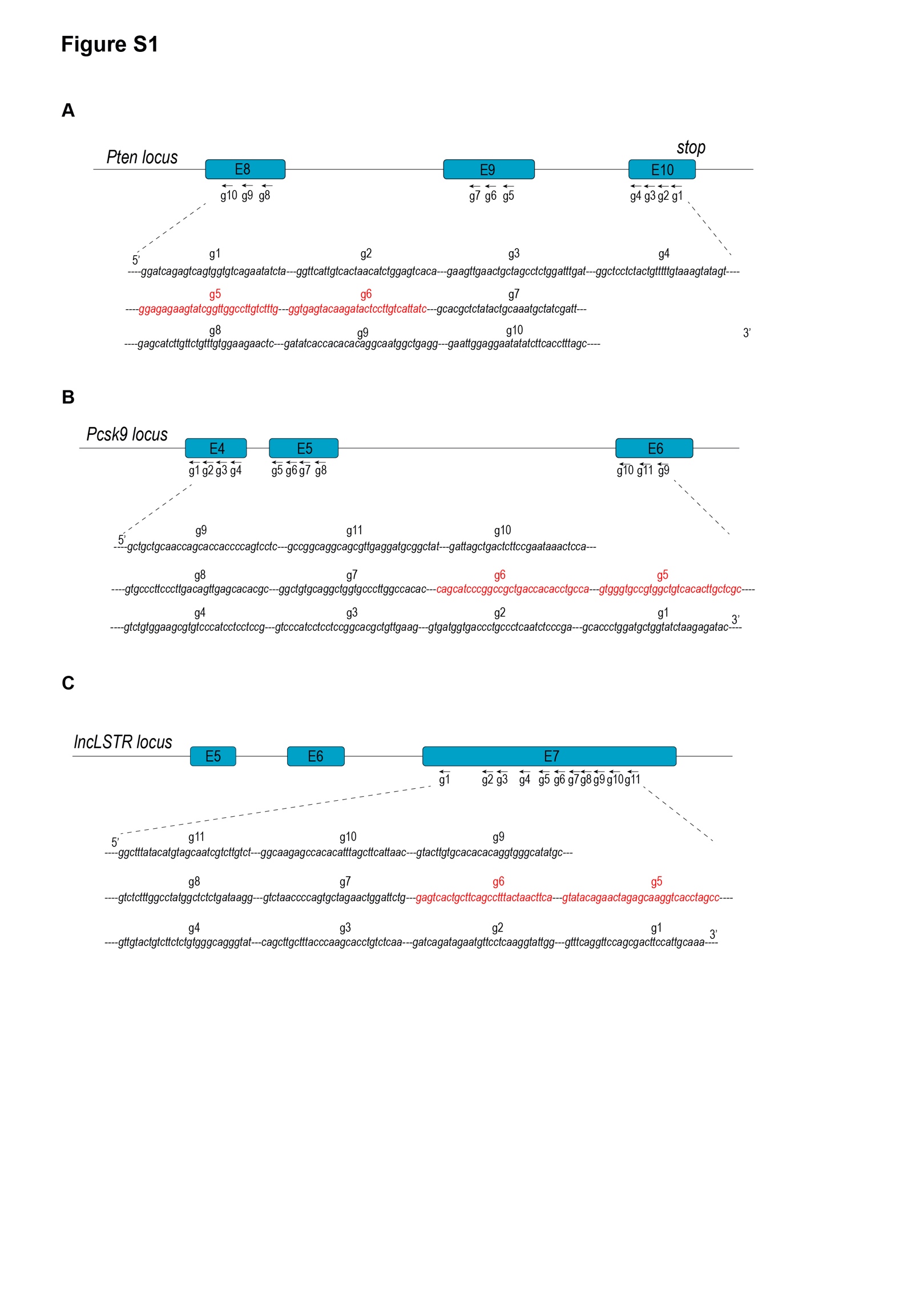

**Fig. S1: Design of sgRNAs.**

**A.** 10 sgRNAs designed in coding sequence of Pten. sgPten-5 (g5, red) and sgPten-6 (g6, red) were selected for in vivo targeted knockdown of Pten.

**B.** 11 sgRNAs designed in coding sequence of Pcsk9. sgPcsk9-5 (g5, red) and sgPcsk9-6 (g6, red) were selected for in vivo targeted knockdown of Pcsk9.

**C.** 11 sgRNAs designed in lncLSTR locus for screening. sglncLstr-5 (g5, red) and sglncLstr-6 (g6, red) were selected for in vivo targeted knockdown of lncLSTR.

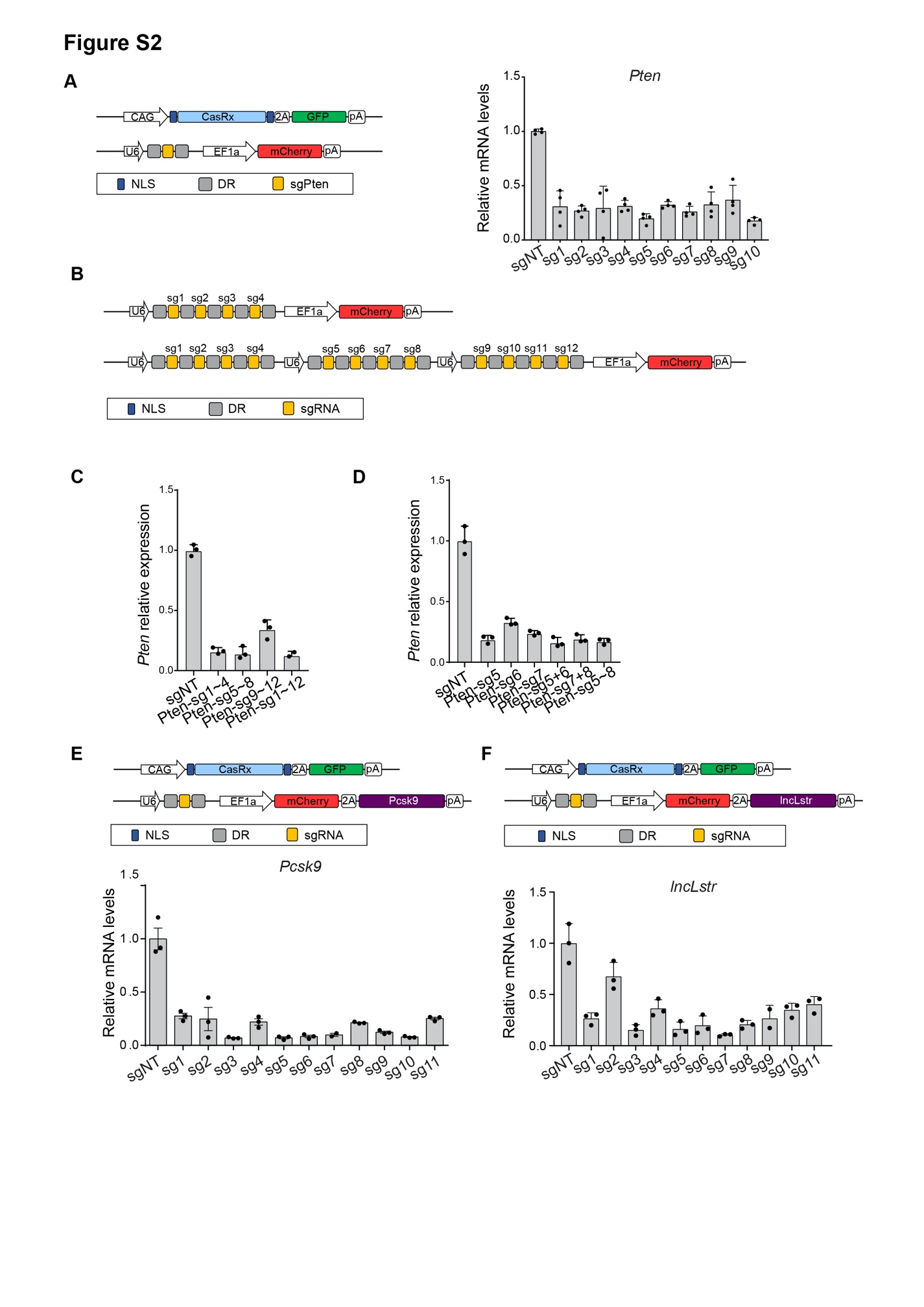

**Fig. S2: CasRx-mediated knock-down of metabolism regulator genes *in vitro***

**A.** N2a cells were transfected with plasmids expressing CasRx and sgRNAs. Left, design of plasmids used for *Pten* sgRNA screen. Right, mRNA levels of *Pten* were quantified in N2a cells receiving CasRx and indicated sgRNA (n=4).

**B.** Schematic of plasmids used for CasRx-mediated *Pten* knockdown in N2a cells with the combination of different sgRNAs.

**C, D.** Knockdown of *Pten* by combination of different sgRNAs in N2a cells (n=3).

**E, F.** 293T cells were transfected with plasmids expressing CasRx, sgRNA and *Pcsk9* (**E**) or *lncLstr* (**F**). Knockdown efficiency of *Pcsk9* (**E**) or *lncLstr* (**F**) was quantified in 293T cells receiving indicated sgRNA (n=3, except for sglncLstr-9, n=2). Data are represented as mean with SD. **p<0.01.

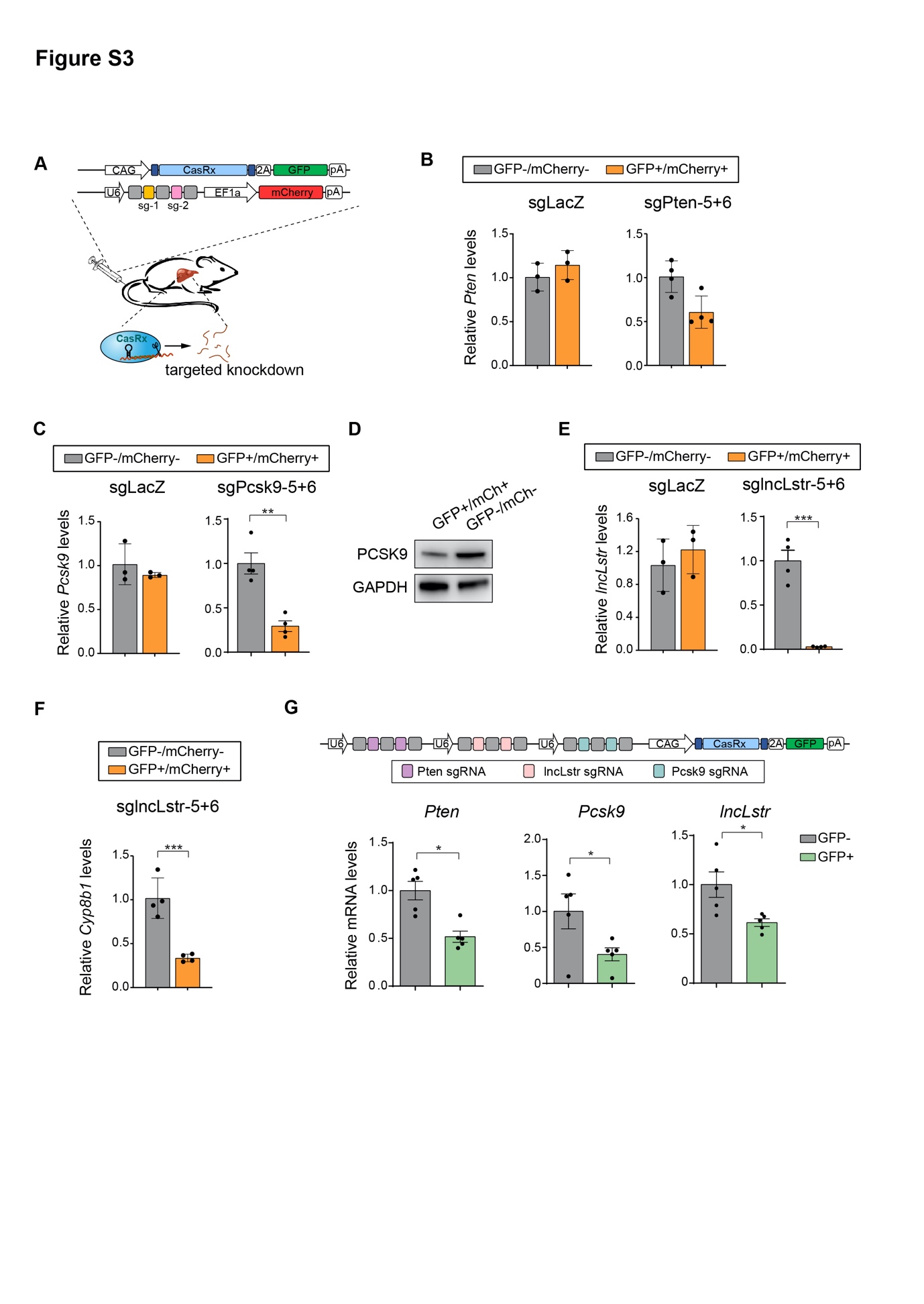

**Figure S3: CasRx-mediated knockdown of metabolic genes in hepatocytes *in vivo***

**A.** Plasmids expressing CasRx and sgRNAs were delivered to wild-type mouse livers by hydrodynamic tail-vein injection.

**B.** Hepatocytes receiving the plasmids encoding CasRx, sgPten-5 and sgPten-6 (GFP+/mCherry+) were purified to quantify the expression of *Pten* by qPCR (n=4). The expressions of *Pten* were not significantly changed in hepatocytes receiving CasRx and sgLacZ (n=3).

**C, D.** CasRx-mediated knockdown of *Pcsk9* in hepatocytes. *Pcsk9* mRNA levels and protein levels were quantified by qPCR (n=4) (**C**) and western blot (**D**). The expressions of *Pcsk9* were not significantly changed in hepatocytes receiving CasRx and sgLacZ (n=3) (**C**).

**E, F.** CasRx-mediated knockdown of *lncLstr* in hepatocytes. *lncLstr* (**E**) and its downstream gene *Cyp8b1* (**F**) were quantified by qPCR (n=4). The expressions of *lncLstr* were not significantly changed in hepatocytes receiving CasRx and sgLacZ (n=3) (**E**).

**G.** CasRx with sgRNA arrays simultaneously knocked down *Pten*, *Pcsk9* and *lncLstr* in hepatocytes (n=5). Data are represented as mean with SD. *p<0.05, **p<0.01, ***p<0.001.

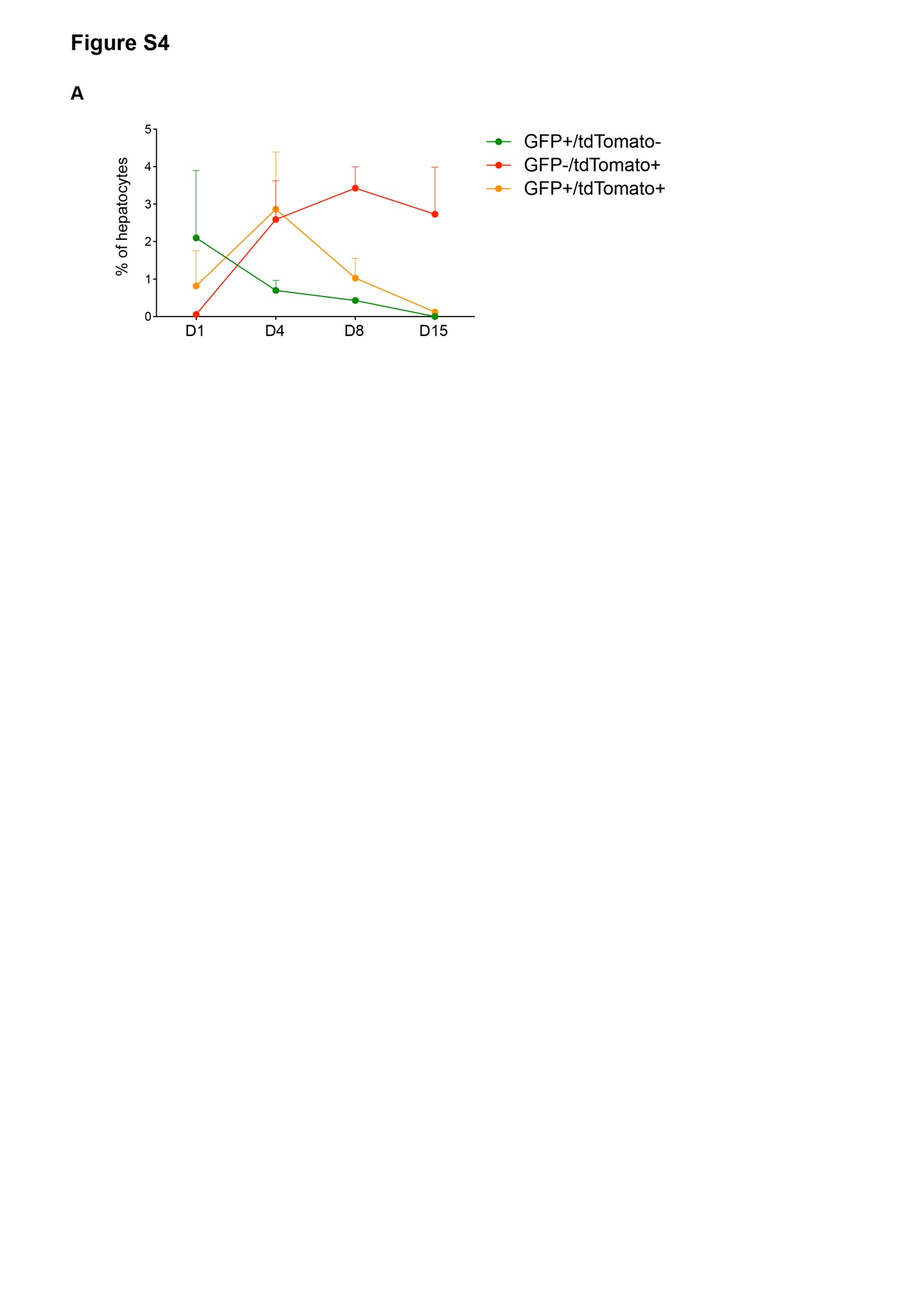

**Fig. S4: Gradually loss of CasRx plasmids in hepatocytes.**

A. The proportions of hepatocytes expressing indicated fluorescence proteins were quantified by flow cytometry at day 1, 4, 8 and 15 after hydrodynamic tail-vein injection of CasRx plasmids into ROSA-CAG-lsl-tdTomato mice (D1, n=4; D4, D8, D15, n=3). Data are represented as mean with SD.

| **Table S1.sgRNA sequences** | |
| --- | --- |
| pten sg1 | ggatcagagtcagtggtgtcagaatatcta |
| pten sg2 | ggttcattgtcactaacatctggagtcaca |
| pten sg3 | gaagttgaactgctagcctctggatttgat |
| pten sg4 | ggctcctctactgtttttgtaaagtatagt |
| pten sg5 | ggagagaagtatcggttggccttgtctttg |
| pten sg6 | ggtgagtacaagatactccttgtcattatc |
| pten sg7 | gcacgctctatactgcaaatgctatcgatt |
| pten sg8 | gagcatcttgttctgtttgtggaagaactc |
| pten sg9 | gatatcaccacacacaggcaatggctgagg |
| pten sg10 | gaattggaggaatatatcttcacctttagc |
| pcsk9 sg1 | gcaccctggatgctggtatctaagagatac |
| pcsk9 sg2 | gtgatggtgaccctgccctcaatctcccga |
| pcsk9 sg3 | gtcccatcctcctccggcacgctgttgaag |
| pcsk9 sg4 | gtctgtggaagcgtgtcccatcctcctccg |
| pcsk9 sg5 | gtgggtgccgtggctgtcacacttgctcgc |
| pcsk9 sg6 | cagcatcccggccgctgaccacacctgcca |
| pcsk9 sg7 | ggctgtgcaggctggtgcccttggccacac |
| pcsk9 sg8 | gtgcccttcccttgacagttgagcacacgc |
| pcsk9 sg9 | ggagtagaggcaggcgtcgtcccggaagtt |
| pcsk9 sg10 | gattagctgactcttccgaataaactcca |
| pcsk9 sg11 | gccggcaggcagcgttgaggatgcggctat |
| lstr sg1 | gtttcaggttccagcgacttccattgcaaa |
| lstr sg2 | gatcagatagaatgttcctcaaggtattgg |
| lstr sg3 | cagcttgctttacccaagcacctgtctcaa |
| lstr sg4 | gttgtactgtcttctctgtgggcagggtat |
| lstr sg5 | gtatacagaactagagcaaggtcacctagcc |
| lstr sg6 | gagtcactgcttcagcctttactaacttca |
| lstr sg7 | gtctaaccccagtgctagaactggattctg |
| lstr sg8 | gtctctttggcctatggctctctgataagg |
| lstr sg9 | gtacttgtgcacacacaggtgggcatatgc |
| lstr sg10 | ggcaagagccacacatttagcttcattaac |
| lstr sg11 | ggctttatacatgtagcaatcgtcttgtct |
| lacz sg | cgtctggccttcctgtagccagctttcatc |
| shPTEN-5 | acaaggccaaccgatacttctCTCGAGagaagtatcggttggccttgtTTTTTG |
| shPTEN-6 | atgacaaggagtatcttgtacCTCGAGgtacaagatactccttgtcatTTTTTG |

| **Table S2**. **Primers used for plasmids construction** | |
| --- | --- |
| Pten sg5+6-R1,R2 | gaagaccccttggataatgacaaggagtatcttgtactcaccgtttcaaacccc |
| LSRT sg5+6 F1 | cacctagcccaagtaaacccctaccaactggtcggggtttgaaacgagtcact |
| LSRT sg5+6 R1,R2 | gaagaccccttgtgaagttagtaaaggctgaagcagtgactcgtttcaaacccc |
| LSRT sg5+6 F2 | gaagacctaaacgtatacagaactagagcaaggtcacctagcccaagtaaaccc |
| pcsk9 sg5+6 F1 | acttgctcgccaagtaaacccctaccaactggtcggggtttgaaaccagcatc |
| pcsk9 sg5+6 R1,R2 | gaagaccccttgtggcaggtgtggtcagcggccgggatgctggtttcaaaccccg |
| pcsk9 sg5+6 F2 | gaagacctaaacgtgggtgccgtggctgtcacacttgctcgccaagtaaacc |
| Pten sg5+6 pcr F1 | caggtctcactagtgagggcctatttcccatgat |
| Pten sg5+6 pcr R1 | caggtctcatcagaaataccgcatcagaattcaaa |
| Lstr sg5+6 pcr F | caggtctcactgagagggcctatttcccatgatt |
| Lstr sg5+6 pcr R | caggtctcacttcaaataccgcatcagaattcaaa |
| 2A-F | gcggcatggacgagctgtacaagggcagtgga |
| 2A-R | cgacgtcaccgcatgttagcagacttcctctgccctctccactgcccttgtacagc |
| Lstr-F | ctaacatgcggtgacgtcgaggagaatcctggcccaagtctgtcaaactgaatggg |
| Lstr-R | taaacaagttttacttgtagcttatcaatccaatgcaccattttactccaatatggt |
| Pcsk9-F | ctaacatgcggtgacgtcgaggagaatcctggcccagatggaagcagccaggtg |
| Pcsk9-R | taaacaagttttacttgtatcactctggagcagaagctgggg |
| pten-sg5+6 pcr F2 | gattcgacattgattattgagatgggctggccttttgctcag |
| pten-sg5+6 pcr R2 | aattgattactattaataactagtcaataatcaatgtccgtaaggagaaaataccgcat |

| **Table S3. qPCR primers** |  |
| --- | --- |
| mouse Pten qPCR F | tggattcgacttagacttgacct |
| mouse Pten qPCR R | gcggtgtcataatgtctctcag |
| mouse Pcsk9 qPCR F | gcccatcgggagattgagg |
| mouse Pcsk9 qPCR R | ttcccttgacagttgagcaca |
| mouse lncLSTR qPCR F | caggtgcttgggtaaagcaa |
| mouse lncLSTR qPCR R | taggaggcagagcttgcttg |
| mouse Gapdh F | ccgtagacaaaatggtgaaggt |
| mouse Gapdh R | cgtgagtggagtcatactggaa |
| human Gapdh F | gtggacctgacctgccgtct |
| human Gapdh R | ggaggagtgggtgtcgctgt |
| mouse Actb qpcr F | gtgacgttgacatccgtaaaga |
| mouse Actb qpcr R | gccggactcatcgtactcc |
| mouse Fasn qpcr F | ggaggtggtgatagccggtat |
| mouse Fasn qpcr R | tgggtaatccatagagcccag |
| mouse Srebf1 qpcr F | gcagccaccatctagcctg |
| mouse Srebf1 qpcr R | cagcagtgagtctgccttgat |
| mouse G6pc qpcr F | cgactcgctatctccaagtga |
| mouse G6pc qpcr R | gggcgttgtccaaacagaat |
| mouse PEPCK qpcr F | ctgcataacggtctggacttc |
| mouse PEPCK qpcr R | gccttccacgaacttcctcac |
| mouse Cyp8b1 qpcr F | cacggggatgtcttcacgg |
| mouse Cyp8b1 qpcr R | tgagcaccagttcttttgcatag |

| **Table S4. Antibodies** |  |  |
| --- | --- | --- |
| anti-PTEN | Cell Signaling | Cat# 9559 |
| anti-pAKT S473 | Cell Signaling | Cat# 4060 |
| anti-AKT | Cell Signaling | Cat# 9272 |
| anti-GAPDH | Proteintech | Cat# 10494-1-AP |
| PCSK9 (human) Polyclonal Antibody | Cayman | Cat# 10007185 |
| HRP-labeled Goat Anti-Rabbit IgG (H+L) | Beyotime | Cat# A0208 |
| Cy3-conjugated AffiniPure Donkey Anti-Rabbit IgG (H+L) | Jackson | Cat# 711-165-152 |

| **Table S5. Software and Algorithms** | | |
| --- | --- | --- |
| SolexaQA v3.1.7.1 | Murray P Cox et al.,2010 | <https://sourceforge.net/projects/solexaqa/> |
| hisat2 v2.0.4 | Daehwan Kim et al.,2015 | <https://ccb.jhu.edu/software/hisat2/index.shtml> |
| Deseq2 1.24.0 | Michael I Love et al.,2014 | <https://bioconductor.org/packages/release/bioc/html/DESeq2.html> |
| Htseq v0.11.2 | Simon Anders et al.,2015 | <https://htseq.readthedocs.io/en/release_0.11.1/count.html> |
